# Modelling the modulation of cortical Up-Down state switching by astrocytes

**DOI:** 10.1101/2022.03.10.483735

**Authors:** Lisa Blum Moyse, Hugues Berry

## Abstract

Converging experimental reports have shown that the firing dynamics of neural networks in several cortical brain areas can exhibit Up-Down synchronization regimes, spontaneously alternating between long periods of high collective firing activity (Up state) and long periods of relative silence (Down state). The molecular or cellular mechanisms that support the emergence of these reversible transitions are still uncertain. In addition to intrinsic mechanisms supported by the local neurons of the network, recent experimental studies have suggested that the astrocytes of the local network can actually control the emergence of Up-Down regimes. Here we propose and study a neural network model to explore the implication of astrocytes in this dynamical phenomenon. We consider three populations of cells: excitatory neurons, inhibitory neurons and astrocytes, interconnected by gliotransmission events, from neurons to astrocytes and back. We derive two models for this three-population system: a rate model and a stochastic spiking neural network with thousands of neurons and astrocytes. In numerical simulations of these three-population models, the presence of astrocytes is indeed observed to promote the emergence of Up-Down regimes with realistic characteristics. Linear stability analysis reveals that astrocytes in these models do not change the bifurcation structure of these systems, but change the localization of the bifurcations in the parameter space. Accordingly, with the addition of astrocytes, the network can enter a bistability region of the dynamics, where the Up-Down dynamical regime emerges. Simulations of the stochastic network model further evidence that astrocytes provide a stationary and stable background of gliotransmission events to the neurons, that triggers spontaneous transitions between synchronized Up and Down phases of neuronal firing. Taken together, our work provides a theoretical framework to test scenarios and hypotheses on the modulation of Up-Down dynamics by gliotransmission from astrocytes.

## 1 Introduction

Collective behaviors, i.e. the emergence of coherent group dynamics from simple interacting elements, are ubiquitous in nature and observed at all scales of biology, from cells to animal populations. Understanding the relationship between the properties of individual elements and a coordinated behavior at the population level usually demands theoretical approaches, for instance from theoretical physics (e.g. Chaté et al., 2008; Hakim and Silberzan, 2017; Cavagna et al., 2018). Among the numerous forms of collective behaviors reported in the brain, Up-Down dynamics characterize a regime of firing dynamics whereby neural networks in several cortical areas exhibit spontaneous transitions between periods of strong collective firing (Up state) and periods of silence (Down state), even in the absence of external inputs (Steriade et al., 1993; Cowan and Wilson, 1994; Cossart et al., 2003; Shu et al., 2003).

The cellular and network mechanisms at the origin of cortical Up-Down dynamics are still not well understood. For a large part, the phenomenon seems intrinsic to the cortical networks since it has been observed in cortical slices (Cossart et al., 2003) and survives *in vivo* when the connections between cortex and thalamus are lesioned (Steriade et al., 1993). A number of theoretical studies have proposed intrinsic mechanisms to explain cortical Up-Down dynamics (e.g. Bazhenov et al., 2002; Compte et al., 2003; Hill and Tononi, 2005; Benita et al., 2012). These proposals usually postulate some sort of activity-dependent negative feedback of the firing rate, according to which individual neurons tend to decrease their firing rate after sustained periods of firing, and to increase it after sustained periods of silence. In the simplest cases, this negative feedback can rely on a slow adaptation current (Compte et al., 2003) or short term plasticity (Hill and Tononi, 2005; Benita et al., 2012), for instance. The existence of a local rhythm generation mechanism intrinsic to the local network does not preclude the impact of input from other brain regions. In particular oscillatory inputs from the thalamus have been suggested to strongly impact or even trigger cortical Up-Down dynamics (Rigas and Castro-Alamancos, 2007; David et al., 2013; Lemieux et al., 2014). Accordingly, several theoretical studies have been proposed to study Up-Down dynamics in the framework of the interplay between an intrinsic activity-dependent negative feedback of the firing rate and an external input to the network (Holcman and Tsodyks, 2006; Lim and Rinzel, 2010; Jercog et al., 2017).

Recently, astrocytes have been identified by experimental studies as a new potential actor of population oscillations in the brain (Lee et al., 2014; Bellot-Saez et al., 2018; Buskila et al., 2019). Astrocytes are non-neuronal neural cells that, together with oligodendrocytes, ependymal cells and microglia form the glial cells (for reviews see e.g. Jkel and Dimou, 2017; Verkhratsky and Nedergaard, 2018). Astrocytes can ensheath synaptic elements, thus forming a “tripartite” synapse where signalling information can flow between the presynaptic neuron, the postsynaptic neuron and the astrocyte (Perea et al., 2009; Araque et al., 2014). Indeed, at the tripartite synapse, astrocytes integrate neuronal activity as a complex transient signal of their intracellular Ca^2+^ concentration (Rusakov, 2015; Shigetomi et al., 2016). In addition, astrocytic intracellular Ca^2+^ signals can, at least under certain conditions, trigger the release by the astrocyte of neuroactive molecules called gliotransmitters that may in turn modulate neuronal information transfer (Santello et al., 2019; Noriega-Prieto and Araque, 2021). The existence in physiological conditions of such a bilateral signalling between neurons and astrocytes is still debated among experimental neuroscientists, in particular regarding the impact of gliotransmitters on neurons (see e.g. Savtchouk and Volterra, 2018; Fiacco and McCarthy, 2018). But if confirmed, it could explain the accumulated experimental evidence of the implication of astrocytes in information treatment in the brain (reviewed in Oliveira et al., 2015; Guerra-Gomes et al., 2017; Santello et al., 2019).

A series of experimental studies by K. Poskanzer and R. Yuste has suggested the implication of astrocytes in the intrinsic mechanism that generates Up-Down dynamics in cortical networks (Poskanzer and Yuste, 2011, 2016). In cortical slices, they observed that increasing calcium activity of a single astrocyte is enough to roughly double the probability to observe an Up state, with no change of the amplitude nor the duration of these Up states (Poskanzer and Yuste, 2011). In vivo experiments further showed that increasing calcium activity in a local population of astrocytes was temporally correlated to a shift of the local neural network to the Up-Down regime (Poskanzer and Yuste, 2016). However, the mechanism by which astrocytes modulate Up-Down cortical dynamics is still unknown. In particular, it is not understood how the modulation by astrocytes interact or rely on the other identified mechanisms of Up-Down state generation.

Here, we used mathematical modeling to propose elements of answers to these questions. We started from the model proposed by Jercog et al. (2017) to model Up-Down cortical dynamics and extended it with the addition of the effect of a population of astrocytes. We expressed the resulting system both as a rate model and an equivalent stochastic spiking neural network. Numerical simulations of both models confirm that astrocytes are indeed potential modulators of Up-Down dynamics. Stability analysis shows that astrocytes do not change the bifurcation structure of these systems, but alter the parameter values for which Up-Down dynamics are likely. As a result, the presence of the astrocyte population can induce the emergence of Up-Down dynamics in a network for which these dynamics are not observed in the absence of astrocytes (at constant parameter values). Interestingly, the difference of signalling timescales between astrocytes and neurons (seconds versus milliseconds) explains a dynamical regime where the astrocytes provide a stationary and stable background of gliotransmission events to the neurons. This background input from the astrocytes, though loosely correlated to the neuronal firing phases, sustains spontaneous transitions between synchronized Up and Down phases of neuronal firing.

## 2 Methods

### 2.1 Rate model

The model of Jercog et al. (2017) was designed to study the emergence of Up-Down dynamics in a neural network composed of an excitatory population E connected to an inhibitory population I in a all-to-all manner (figure 1). The excitatory population is endowed with an adaptation mechanism a, that implements an additive hyperpolarizing current to the population E and that grows with its firing rate. Adaptation is therefore the main intrinsic mechanism for the emergence of Up-Down dynamics in the model. However, each population also receives a fluctuating external input. The firing rate dynamics of this model is given by

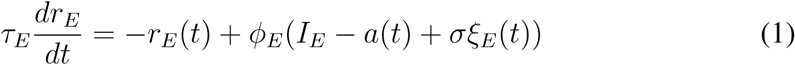

and

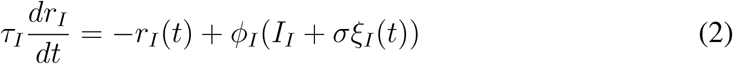

where *r*_*X*_ is the firing rate of population *X* = {*E, I*}, *I*_*X*_ its recurrent inputs (from the populations of the system, see below), and *ξ*_*X*_ a fluctuating external input on *X*, modeled as an independent Ornstein-Uhlenbeck process (see Jercog et al. (2017) for details). *τ*_*X*_ is the time constant of population *X*, and its transfer function:

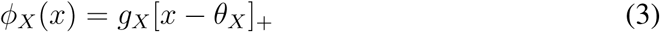

with rectification [*z*]_+_ = *z* if *z* > 0 and 0 otherwise. The dynamics of the adaptation current *a*(*t*) is given by:

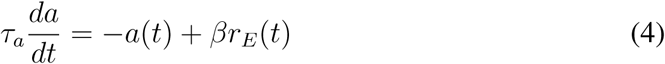

**Figure 1:**
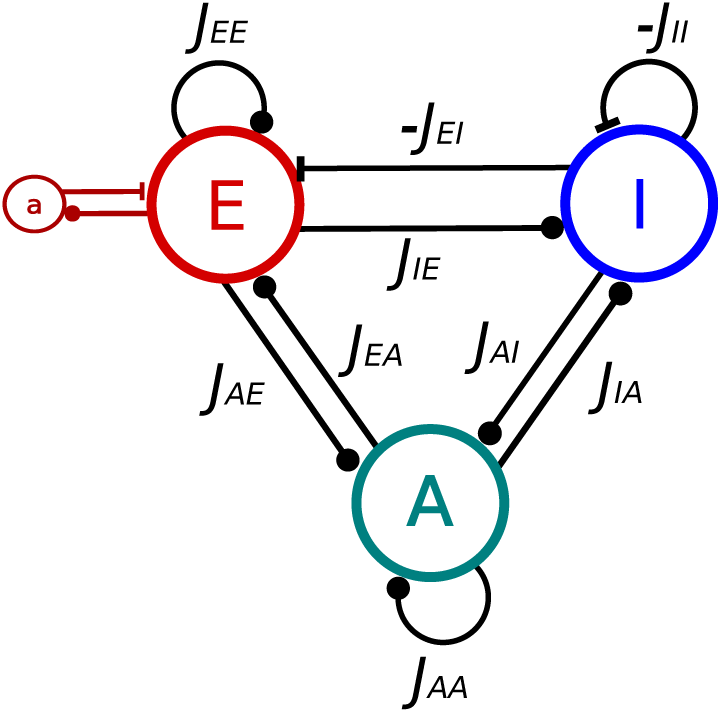
Interactions between the three populations of the model, E for excitatory neurons, I for inhibitory neurons and A for astrocytes. *a* represents the adaptation mechanism of E cells. Lines terminated with a full circle represent positive interactions whereas those terminated with a bar represent inhibition (of I cells on E and I, and adaptation *a* on E cells). *J*_*XY*_ represent the synaptic strength from the *Y* population toward the *X* population. In case of inhibition these terms are negative.

Equations (1) to (4) define the model proposed in Jercog et al. (2017). Here, we extended it to account for the impact of astrocytes on the network.

Astrocytes express a variety of receptors at their membranes, that bind the neurotransmitters or neuromodulators released by the presynaptic elements at the tripartite synapse, including glutamate, GABA, acetylcholine or dopamine (Perea et al., 2009; Verkhratsky and Nedergaard, 2018; De Pitta and Berry, 2019). Through these receptors, neuronal activity is integrated inside the astrocyte as a complex signal of intracellular Ca^2+^ (Rusakov, 2015; Shigetomi et al., 2016). In response to this Ca^2+^ transient, astrocytes can, at least under certain conditions, release in the synapse a variety of molecules, referred to as “gliotransmitters” that, upon binding to the pre- or post-synaptic element of the tripartite synapse will in turn hyperpolarize or depolarize the neuronal membrane potential (Savtchouk and Volterra, 2018; Santello et al., 2019; Noriega-Prieto and Araque, 2021). Interestingly, whereas the astrocytic cytosolic calcium transients are very slow events, especially in the soma (around 10-20 sec on average), gliotransmitter release events are much faster (around 1 sec) (Poskanzer and Yuste, 2016; Bindocci et al., 2017).

According to this oversimplifying birds-eye view of neuron-astrocyte interactions, the astrocytic response to presynaptic neuronal activity is reminiscent of neuronal integration: presynaptic neuronal activity is integrated in astrocytes as a calcium trace that triggers a peak-like release of gliotransmitters that in turn affects postsynaptic membrane voltage. The major differences are i) integration time-scales and gliotransmitter release dynamics in astrocytes are different from electrical signalling in neurons and ii) the equivalent of inhibitory/depolarizing neuronal inputs that decrease the membrane potential does not seem to exist for astrocytic Ca^2+^. To model astrocyte activity, we thus opted for same formalism of rate equations as eq. (1) or eq. (2), but with different time scales and expressed the rate of gliotransmitter release by the astrocyte as

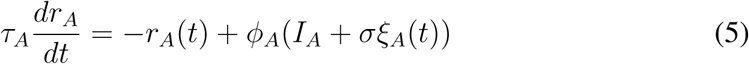

with the constraint *τ*_*A*_ ≫ *τ*_*I*_ and *τ*_*A*_ ≫ *τ*_*E*_. Moreover, whereas one expects positive values for the firing threshold *θ*_*X*_ of neurons in eq. (3) (i.e. neurons remain silent below a threshold of their input), we will favor negative values for *θ*_*A*_, in order to account for the spontaneous calcium activity of astrocytes (Verkhratsky and Nedergaard, 2018).

We now can give a definition for the three internal recurrent inputs:

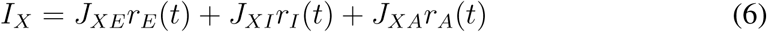

with now *X* = {*E, I, A*}. The synaptic couplings *J*_*XY*_ (with *X, Y* = {*E, I, A*}), describe the strength of the connection from population Y to X. They verify *J*_*XE*_ > 0 (excitatory), *J*_*EI*_ < 0, *J*_*II*_ < 0 (inhibitory) and *J*_*AX*_ ≥ 0 (i.e. both E and I increase the rate of gliotransmitter release in astrocytes).

A fixed-point and linear stability analysis of the rate model defined by equations (1) to (6) is provided in section A of S1 Text. The values of the parameters in the equations above are given in table 1.

**Table 1:**
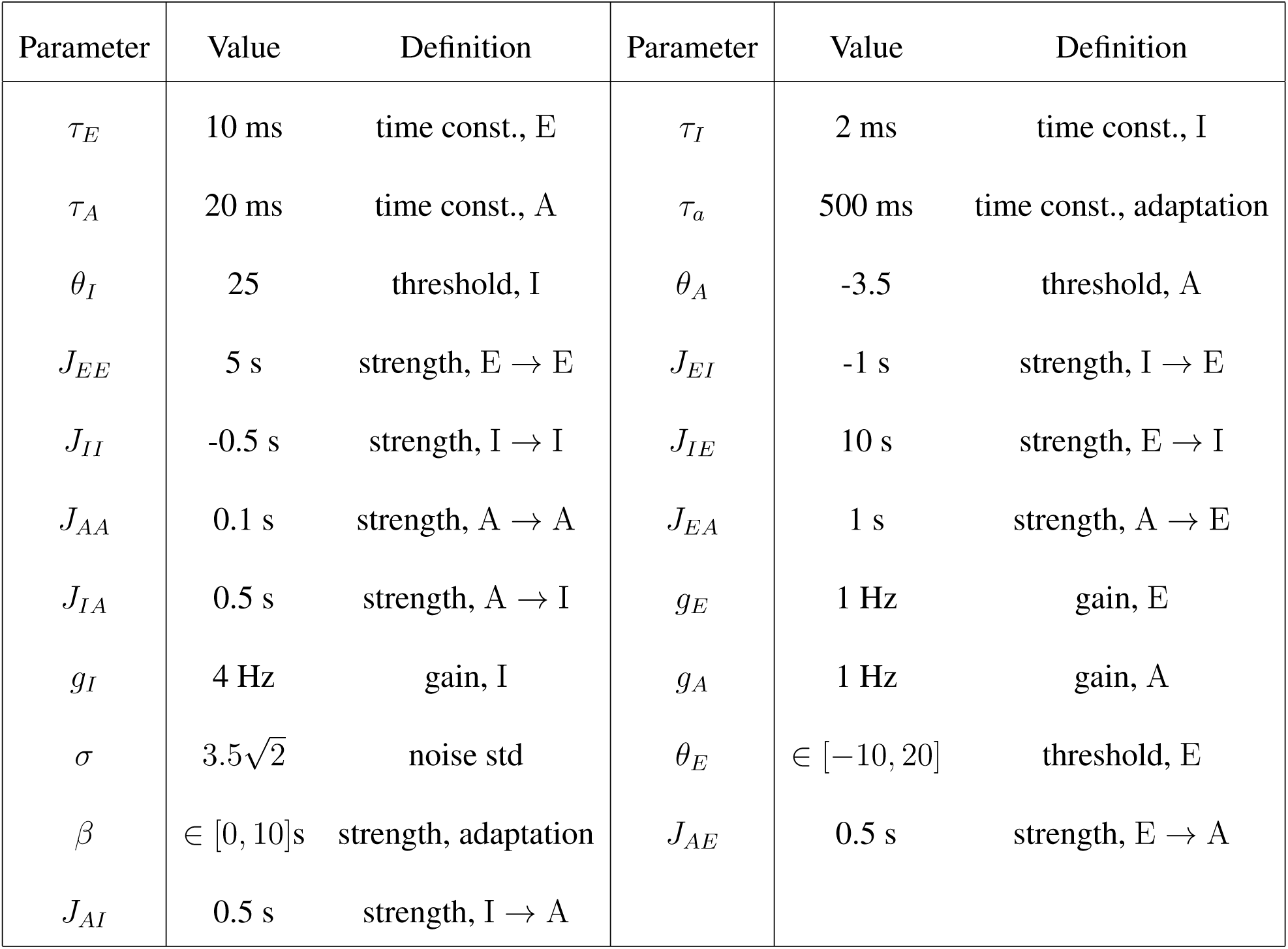
Parameters used for the rate model equations (1) to (6)

### 2.2 Spiking model

We also modeled the three-population {*E, I, A*} system of figure 1 by expressing it as a stochastic spiking network model instead of the firing rate framework of section 2.1. Following the same principle as above, where we used a classical neuron rate equation to model astrocyte gliotransmitter release, we used here leaky integrate and fire equations to model both neuronal membrane potential and the release of gliotransmitter by astrocyte. Hence, the membrane potential of the two populations of neurons reads:

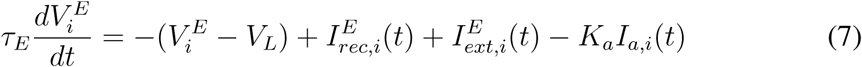

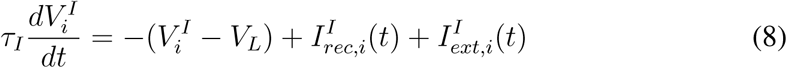

and similarly, we model gliotransmitter release from the astrocytes as:

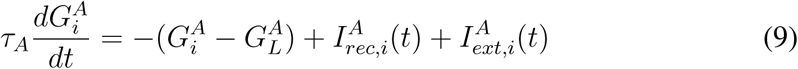

with *i* ∈ {1, …,*N*_*X*_}. 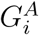 is thus a phenomenological dimensionless variable that integrates the neuronal and astrocytic inputs to astrocyte *i*. According to the integrate-and-fire principle, whenever the membrane voltage of a neuron of population X exceeds its threshold *θ*_*X*_ at time *t*, a spike is emitted and the membrane voltage is reset to 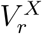. Similarly, when 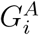 exceeds the threshold *G*_*th*_, astrocyte *i* emits a gliotransmitter release event, and 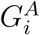 resets to *G*_*r*_. Gliotransmitter release events and spikes are then integrated in the corresponding synaptic variable *s*_*X*_ (see equations (11) and (12) below).

For the simulations of the spiking model, we assumed the following connectivity rules:

- full connectivity for neuron-to-neuron connections and for astrocyte-to-astrocyte connections. The latter emulates the organization of astrocytes as a syncytium (Verkhratsky and Nedergaard, 2018).
- only a fraction (10%) of the E or I neurons are subjected to gliotransmission from the astrocytes. These neurons are chosen at random (uniform distribution).
- one half of the astrocytes, chosen uniformly at random, receive inputs from the E or I neurons

One specificity of the original spiking model of Jercog et al. (2017) is to account for synaptic variables by a single pair of variables *u*_*X*_, *s*_*X*_ for each population, which can thus be considered as population variables instead of individual cell variables. Here we follow this model and define the recurrent input to each population *X* = {*E, I, A*} as a population-level input as:

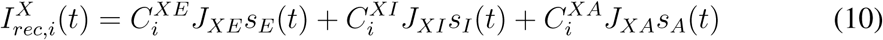

where the A → E connectivity 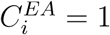 for 10% of the E neurons i (chosen uniformly at random) and 0 for the others (the same for 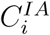). For the (E, I) → A connectivity, we used 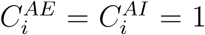 for 50% of the astrocytes (chosen uniformly at random) and 0 for the others. All the others connectivities 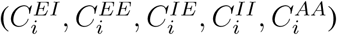 were set to 1 (all-to-all connectivity).

The synaptic variables *s*_*X*_ integrate the spikes or release events emitted by all the cells in population X:

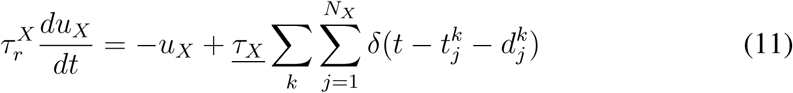

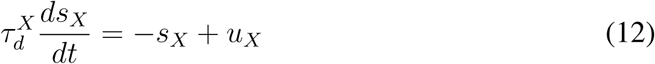

with 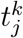 the *k*^th^ spike (or release) time of cell *j* of population X, 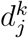 its transmission delay (uniformly distributed between 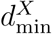 and 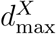), and 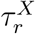 and 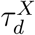 the rise and decay times of the synapse, respectively.

Note that signal transmission in astrocytes is much slower than in neurons since it is based on reaction-diffusion (calcium signalling) instead of the propagation of an action potential (Poskanzer and Yuste, 2016; Bindocci et al., 2017; Verkhratsky and Nedergaard, 2018). To account for this important difference in timescales, we used transmission delays that were on the order of milliseconds for neurons ([0, 1] ms) but on the order of seconds for astrocytes ([0.5, 1.5] s, see table 2).

**Table 2:** Parameters used for the spiking model equations (7) to (13)

| Parameter | Value | Definition | Parameter | Value | Definition |
| --- | --- | --- | --- | --- | --- |
| $\tau_E$ | 20 ms | time const., E | $\tau_I$ | 10 ms | time const., I |
| $\tau_A$ | 160 ms | time const., A | $\tau_a$ | 500 ms | time const., adaptation |
| $\underline{\tau_E}$ | 1 ms | time const., $u_E$ | $\underline{\tau_I}$ | 1 ms | time const., $u_I$ |
| $\underline{\tau_A}$ | 1 ms | time const., $u_A$ | $J_{EE}$ | 1.4 mV | strength, E → E |
| $J_{EI}$ | -1.4 mV | strength, I → E | $J_{II}$ | -1 mV | strength, I → I |
| $J_{IE}$ | 1.25 mV | strength, E → I | $J_{AA}$ | 0.16 | strength, A → A |
| $J_{AE}$ | 0.053 | strength, E → A | $J_{EA}$ | 22 mV | strength, A → E |
| $J_{IA}$ | 4.4 mV | strength, A → I | $J_{AI}$ | 0.058 | strength, I → A |
| $\beta$ | 1 ms | time const., adaptation | $K_a$ | 600 | strength, adaptation |
| $\sigma_E$ | 3 mV | noise std, E | $\sigma_I$ | 3 mV | noise std, I |
| $\sigma_A$ | 3 | noise std, A | $V_r$ | 14 mV | reset membr. pot. |
| $G_r$ | 9 | reset gliotrans. release | $V_{th}$ | 20 mV | spike-threshold |
| $G_{th}$ | 13 | gliotrans. release thresh. | $\tau_d^E$ | 23 ms | decay time, E |
| $\tau_d^I$ | 1 ms | decay time I | $\tau_d^A$ | 2 ms | decay time, A |
| $\tau_r^E$ | 8 ms | rise time, E | $\tau_r^I$ | 1 ms | rise time, I |
| $\tau_r^A$ | 8 ms | rise time, A | $d_{\min}^E$ | 0 ms | min. delay, E |
| $d_{\min}^I$ | 0 ms | min. delay, I | $d_{\max}^E$ | 1 ms | max. delay, E |
| $d_{\max}^I$ | 0.5 ms | max. delay, I | $d_{\min}^A$ | 500 ms | min. delay, A |
| $d_{\max}^A$ | 1.5 s | max. delay, A | $V_L^E$ | 7.6 mV | leak potential, E |
| $V_L^I$ | 6.5 mV | leak potential, I | $G_L^A$ | 7 | leak gliotrans. rate, A |

In addition, the excitatory neurons displayed an after hyperpolarization (AHP) current:

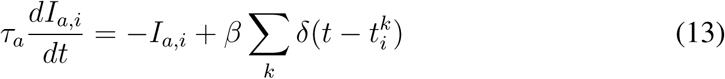

The external input current 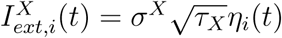 is a Gaussian white noise term.

Initial conditions were set as 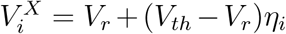 and 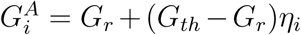, where *η*_*i*_ is a random value with uniform distribution between 0 and 1. Unless indicated, we simulated the spiking network model using *N*_*E*_ = 4,000 excitatory neurons, *N*_*I*_ = 1,000 inhibitory neurons and *N*_*A*_ = 2, 000 astrocytes. Each of these 2,000 astrocytes thus impacts 400 excitatory and 100 inhibitory neurons by gliotransmitter release, whereas half of them are individually impacted by the activity of the totality of the 4,000 E and 1,000 I neurons.

The values of the parameters in the equations above are given in table 2. A fixed-point and linear stability analysis of the spiking model defined by equations (7) to (13) is provided in section B of S1 Text.

### 2.3 Automatic segmentation of Up and Down phases

We quantified the statistics of Up and Down phases in rate model or spiking network simulations based on the mean firing rate time series. In spiking network simulations, we first computed the mean population rate from the raster plot, using a sliding window of 10 ms and counting the total number of spikes emitted by all neurons (excitatory and inhibitory) during the window. Automatic segmentation of the firing rate time series into Up phases and Down phases was achieved by smoothing the sampling rate using a sliding window of +/− 50 points around each data point and replacing each data point by the median over the window. Transition of the smoothed data curve through a threshold of 1.0 Hz from below was considered a switch from a Down to a Up state, whereas transition from above signaled a reverse switch, from Up to Down state. The first and last phases of a simulation were systematically discarded and not accounted for in the statistics.

## 3 Results

### 3.1 Rate model

We first illustrate the dynamics of the rate model described in section 2.1 by the simple numerical simulations of figure 2. In the absence of astrocytes (i.e. with *J*_*IA*_ = *J*_*EA*_ = *J*_*AI*_ = *J*_*AE*_ = 0 s, fig. 2a), the model with the parameters of the figure is silent: the firing rate of the inhibitory neurons *r*_*I*_ vanishes, and that of the excitatory neurons, *r*_*E*_, is also zero most of the time, except for small fluctuations due to external noise. Accordingly, adaptation is essentially off. We then added gliotransmission between excitatory neurons and astrocytes in fig. 2b keeping all other parameters identical to fig. 2a. Adding gliotransmission drastically changes the dynamics that now exhibits spontaneous transitions between long periods of silence for all neuronal populations and shorter periods of high firing rates for the excitatory and inhibitory neurons (around 10 and 5 Hz, respectively). In other words, astrocyte activity in the rate model switches the dynamics from silent to a Up-Down oscillatory dynamics, in agreement with experimental observations *in vivo* (Poskanzer and Yuste, 2016). During an Up state, adaptation slowly increases and eventually triggers the Up-to-Down transition that ends the Up state.

**Figure 2:**
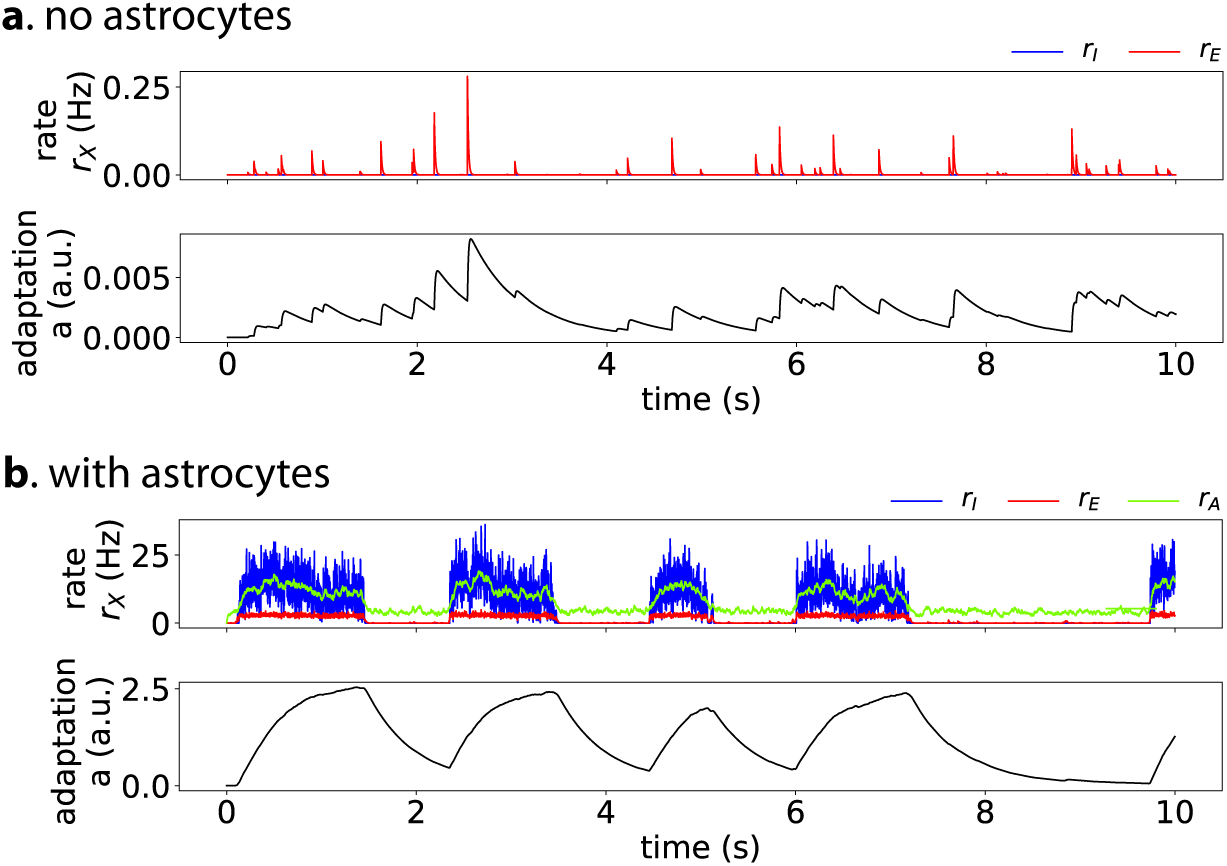
Astrocytes in the rate model of sec. 2.1 switch the dynamics from silent to Up-Down. In the absence of astrocytic impact on the neurons (*J*_*EA*_ = *J*_*AE*_ = *J*_*IA*_ = *J*_*AI*_ = 0 s) **(a)**, the neurons are in a silent state with vanishing firing rates *r*_*E*_ (*red*) and *r*_*I*_ (*blue*) and adaptation *a* (*black*), corresponding to a Down-state fixed point. Please note the difference of *y*-scale between panels (a) and (b). When gliotransmission between astrocytes and neurons is accounted for (**b**, *J*_*EA*_ = 1, *J*_*IA*_ = *J*_*AI*_ = *J*_*AE*_ = 0.5 s), *with no change of the other parameters*, the dynamics switches to Up-Down dynamics. All other parameters given in table 1.

Note that the average values of *r*_*E*_, *r*_*I*_ and *r*_*A*_ during the Up and Down states in the simulation of the figure match the values predicted by the stability analysis of section A of S1 Text. In particular, this analysis states that in the Down fixed-point, one expects *r*_*A*_ = −*g*_*A*_*θ*_*A*_/(1 − *g*_*A*_*J*_*AA*_) while the neuronal rates vanish. In agreement, the rate of gliotransmitter release by the astrocytes *r*_*A*_ remains elevated during the Down states of fig. 2b, even though the neurons are silent.

To analyze further these simulation results, figure 3 summarizes the fixed-point and linear stability analysis of section A of S1 Text in the (*β, θ*_*E*_)-parameter plan. In the absence of noise, fig. 3A, the system behavior is determined by two straight lines: the Down steady-state exists (and is stable) only on the right hand side of the line defined by eq. (S1.5) whereas the Up steady-state exists for the half-plan below the line defined by eq. (S1.14). This defines two regions of mono-stability, one where the Up state is the only fixed-point (U-region) and the other where the Down state in the only one (D-region). The region where both the Up state and the Down state exist is a region of bistability (“Bist.” in the figure) where the dynamics converges to the Up or the Down state depending on the initial conditions. Finally, in the region where neither the Up nor the Down fixed-points exist, the arguments of the rectification functions regularly switch from positive to negative and back. This yields a regime of oscillations (“Osc”-region), that is a specific manifestation of the non-smooth character of the model.

**Figure 3:**
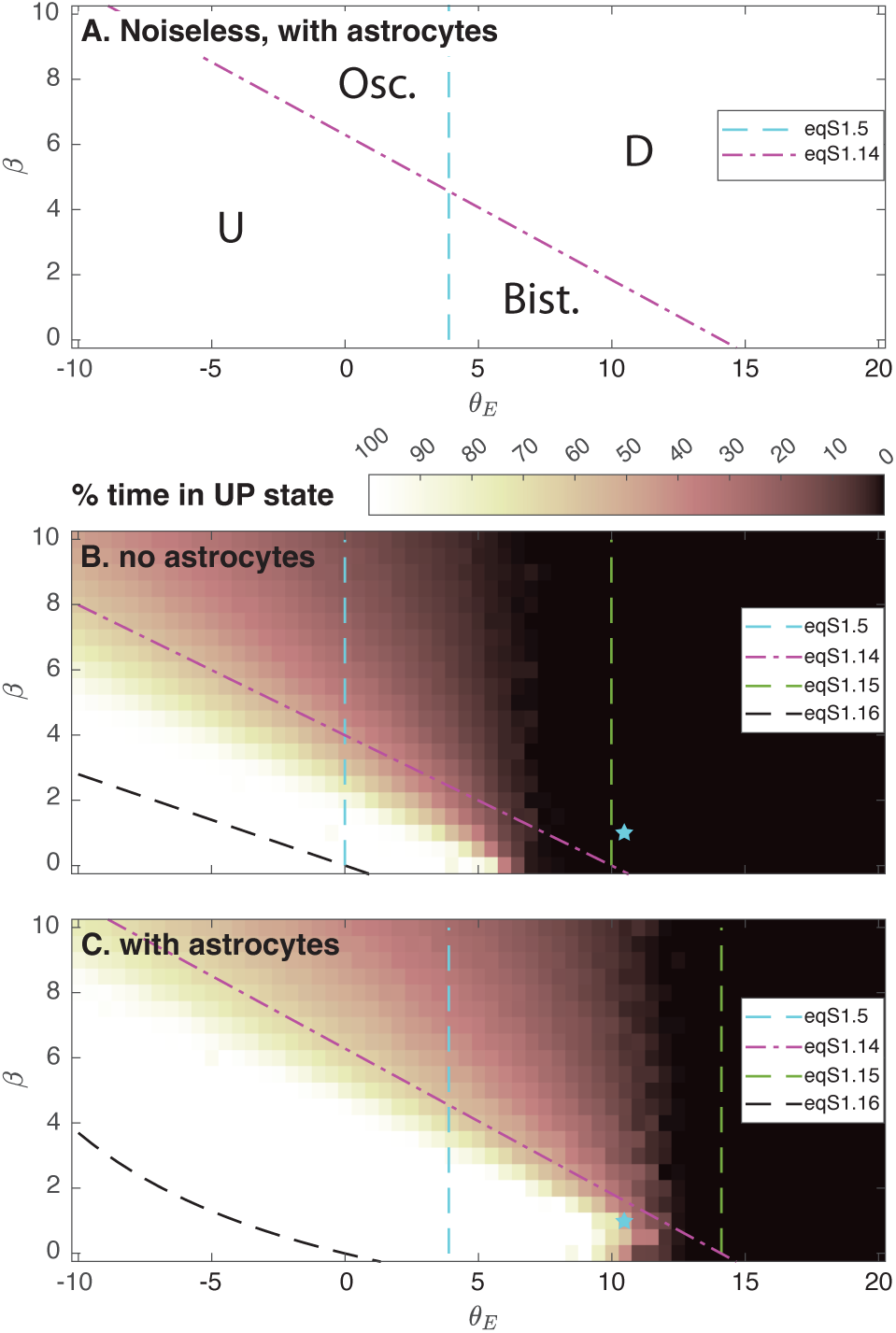
Prediction of the dynamical regimes of the model as a function of the adaptation strength *β* and the threshold of the excitatory neurons *θ*_*E*_. In the absence of noise **(A)**, the deterministic model exhibits two regions of monostability: the “U”-region where the Up state fixed-point is the only one, and the “D”-region where the Down state fixed-point is the unique fixed-point. Both fixed-points coexist in the bistable region “Bist.” whereas the dynamics oscillates in the “Osc.” region. Those regions are precisely delimited by eq. (S1.5) and eq. (S1.14). With noise, three main regimes are predicted: a purely Up state in the bottom left part of the plan, a purely Down state in the right part of the plan and spontaneous transitions between Up and Down states in-between (*U* ↔ *D*). The color-code indicates the percentage of time spent ↔ in the Up state during a simulation. Simulations were carried out with astrocytes (*J*_*EA*_ = 1, *J*_*IA*_ = *J*_*AI*_ = *J*_*AE*_ = 0.5 s, **A, C**) or in their absence (*J*_*IA*_ = *J*_*AI*_ = *J*_*EA*_ = *J*_*AE*_ = 0 s, **B**). The cyan star locates the parameters of figure 2, which shows in particular that gliotransmission pushes the frontiers of the Up-Down region further to the right, effectively switching the dynamics to the Up-Down regime. Equation (S1.15) and eq. (S1.16) are theoretical estimates of the frontiers between *U* ↔ *D* and *D* or *U* ↔ *D* and *U*. All other parameters given in table 1.

With noise (fig. 3B-C), spontaneous Up-Down transitions are expected to occur in the bistable and oscillatory regions of the noiseless system, but also in sub-regions of its U- and D-regions (see S1 Text). Altogether, this defines three dynamical regimes: low values of *β* and *θ*_*E*_ are predicted to give rise to a regime of perpetual high firing rates, i.e. a stable Up state. Conversely, large values of the excitatory neuron threshold *θ*_*E*_ are expected to yield a silent regime, or Down state, where neuronal firing rates vanish. Between those two regions, the system is predicted to switch spontaneously between periods of high population rates and periods or collective silence, i.e. *U* ↔ *D* dynamics. A theoretical estimation for the frontier between the *U* and the *U* ↔ *D* region with noise is given by eq. (S1.16) (fig. 3, *dashed black lines*). Comparing with the percentage of the simulation-time spent in the Up state, it seems that in all the cases illustrated in fig. 3B-C, this expression indeed correctly positions the *D*-region on the plan, although it strongly underestimates its size. Likewise eq. (S1.15) (fig. 3, *dashed green lines*) indeed indicates the transition between the *D* and the *U* ↔ *D* regions, although, here again, the predicted size of the *D* region is strongly underestimated.

Remarkably, this frontier between *U* ↔ *D* and *D* is very sensitive to modifications of gliotransmission couplings (the *J*_*XA*_s and *J*_*AX*_s). As can be seen from the figure, the presence of astrocytes pushes this frontier to larger values of *θ*_*E*_. The cyan star in fig. 3B-C locates the parameter values used in fig. 2. Without astrocytes, the star is located on the right hand side of the frontier between the *U* ↔ *D* and the *D* region, thus explaining the silent state of fig. 2a. With the addition of gliotransmission, the frontier moves rightwards, so that now, the star is located inside the *U* ↔ *D* region. This explains the Up-Down regimes of fig. 2b and c.

Taken together, these results indicate that astrocyte activity can indeed switch the network dynamics from silence to the Up-Down regime by altering the phase diagram of the dynamics. The effect of gliotransmission on the model dynamics is not drastic, in particular gliotransmission does not change the nature or the number of bifurcation points in the system. However, it displaces the frontiers separating dynamical regimes, thus allowing the expression of Up-Down dynamics for a larger range of values of the firing threshold of the excitatory neurons.

### 3.2 Stochastic spiking network model

To assess the significance of the above mechanisms in a more biophysically realistic circuit, we next expressed the circuit of figure 1 as a stochastic spiking network model, with leaky integrate-and-fire neurons and astrocytes (section 2.2). Using the same illustration as for the firing rate model above, we start in figure 4 with a network devoid of astrocytes, i.e. for which *J*_*AE*_ = *J*_*AI*_ = 0 and *J*_*EA*_ = *J*_*IA*_ = 0 mV. The neurons exhibit a very short firing phase at the beginning of the stimulation due to our choice of random initial conditions but quickly converges back to a silent state.

**Figure 4:**
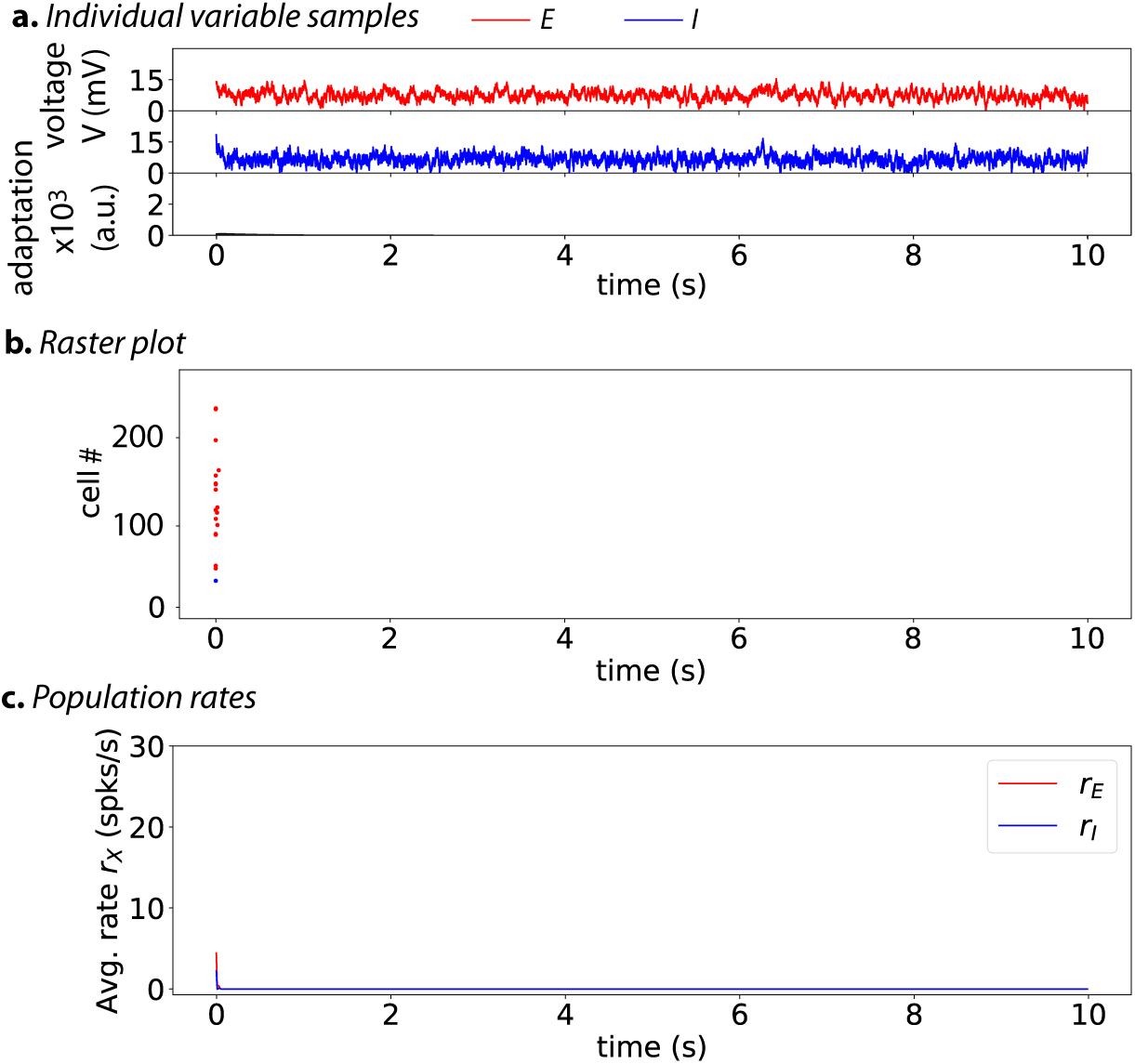
Without astrocytes (*J*_*AE*_ = *J*_*AI*_ = 0, *J*_*IA*_ = *J*_*EA*_ = 0 mV), the stochastic spiking network of sec. 2.2 is in a silent state. The membrane voltages of two randomly chosen cells, one excitatory (*red*) neuron, one inhibitory (*blue*) neuron, as well as the average AHP current (*black*) are shown in **(a)**. The spike rastergram **(b)** that locates with points the spike times of a randomly-chosen subset of the neurons (one neuron = one row), and the corresponding mean population rates **(c)** are shown using the same color-code. The short initial burst of activity is due to the initial conditions where every cell is initiated randomly between its resting potential and the spiking threshold. *N*_*E*_ = 4,000 excitatory neurons, *N*_*I*_ = 1,000 inhibitory neurons. Other parameters given in table 2.

Adding gliotransmission between astrocytes and neurons strongly affects the dynamics (figure 5), yielding spontaneous alternations between periods of nearly complete neuronal silence and periods of high collective neuronal firing, during which roughly all neurons fire on the order of 2 to 15 spikes. The raster plot of fig. 5b also suggests the factors that trigger Down-to-Up transitions: Up states are systematically initiated by a strong firing activity in the subset of excitatory neurons that are contacted by the astrocytes (neurons numbers 50 to 70 in the raster plot). This first wave of excitation then is transmitted to the whole populations of neurons (E and I), thus forming an Up state.

**Figure 5:**
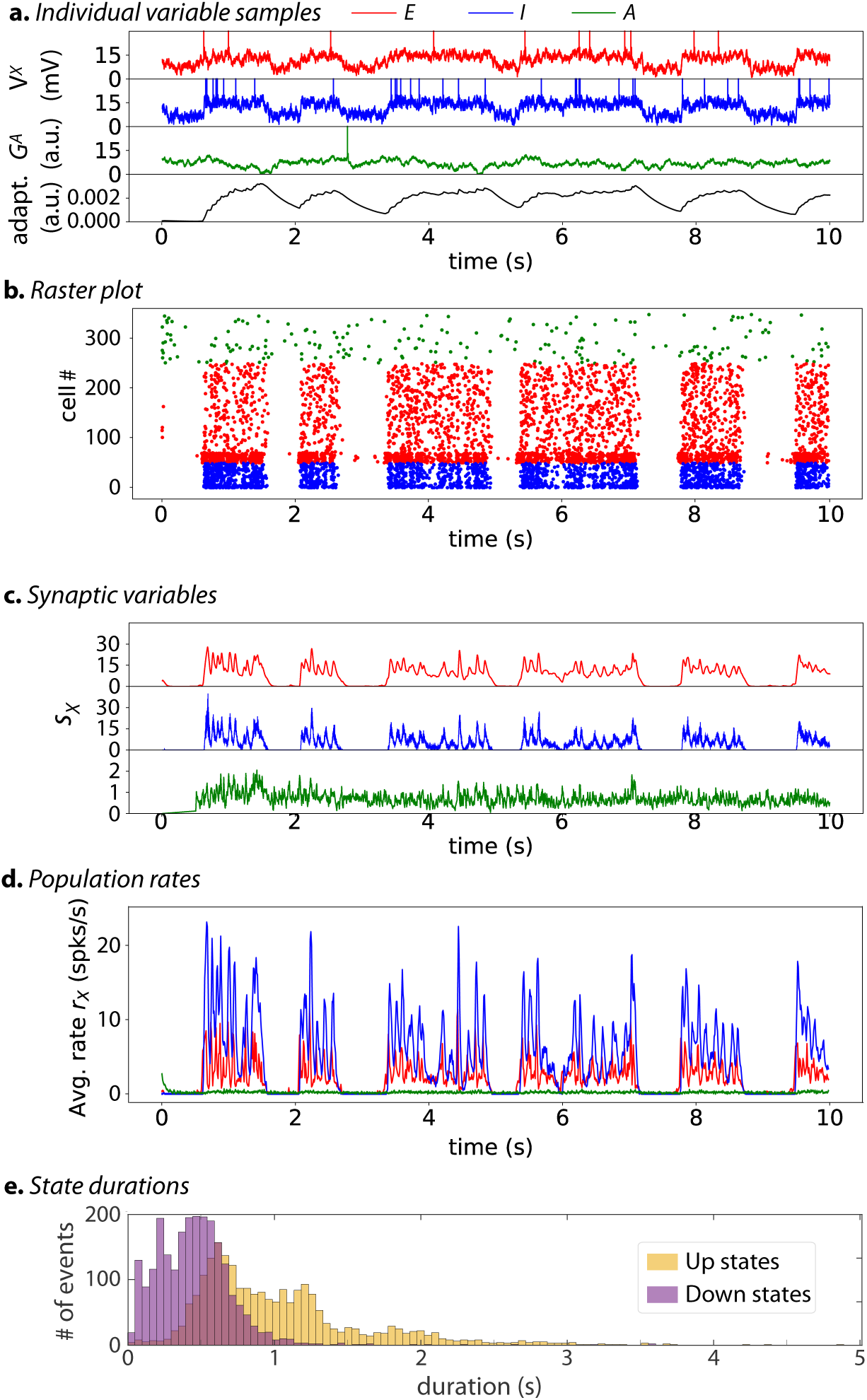
With astrocytes, (*J*_*AE*_ > 0, *J*_*AI*_ > 0), the stochastic spiking network switches to a Up-Down dynamic regime. The membrane voltages of three randomly chosen cells, one excitatory (*red*) neuron, one inhibitory (*blue*) neuron and the astrocyte (*green*) as well as the average AHP current (*black*) are shown in **(a)**. The spike rastergram **(b)**, corresponding synaptic variables *s*_*X*_ **(c)** and mean population rates **(d)** are shown with the same color-code. The distribution of Up (orange) and Down (purple) state durations for E and I cells is shown in **(e)** (based on 200 independent simulations of 20 sec each, resulting in a total of 2273 Up states and 2356 Down). For each simulation, *N*_*E*_ = 4,000 excitatory neurons, *N*_*I*_ = 1,000 inhibitory neurons and *N*_*A*_ = 2, 000 astrocytes. Other parameters given in table 2. For readability, the first phase of the simulation, characterized by a short very active up state, was discarded.

Hence, our biophysical model of stochastic spiking neurons confirms that astrocyte activity can switch the neuronal network from silent to Up-Down dynamics. During the Up states, the mean population firing rate of the spiking network are similar to those exhibited by the firing rate model, i.e. around 10 Hz for inhibitory neurons and 5 Hz for excitatory ones (compare fig. 5c with fig. 2b), confirming the good match between the two models despite the dissimilarity of their spatiotemporal scales. The distributions of the duration of the Up and Down states are estimated in fig. 5d. For Down states the distributions is peaked around 0.5 seconds whereas it is much broader for Up states, with a large part of the durations comprised between 0.5 and 1.3 seconds. On average, the Down states are twice shorter than the Up states: 459 ± 336 ms for the Down states *versus* 1, 031 ± 575 ms for the Up.

However, unlike the neurons that collectively synchronize their firing as successive Up and Down phases, the rate of gliotransmission events by astrocytes does not exhibit strong evidence of alternation between distinct activity phases (fig. 5, *green*). The membrane potential of the individual neurons is strongly bimodal, fluctuating around a lower mean value during Down phases and around a larger mean during Up phases, on top of which spikes are emitted (fig. 5a, *blue, red*). In strong contrast, the dynamics of the gliotransmitter release variables 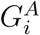 is devoid of such alternations, rather appearing to fluctuate around a single, stationary mean (fig. 5a, *green*). This opposition is also visible in the raster plot (fig. 5b): the neuronal spikes are strongly synchronized and their presence almost totally restricted to the Up phases, whereas the astrocytic gliotransmitter release events are emitted at an intermediate frequency, but with no clear variation of frequency between Up and Down phases. The evolution of the population synaptic variables, the *s*_*X*_s of equations (10) to (12), provides another evidence that the neuronal and astrocytic dynamics are different (fig. 5c): the astrocytic variable *s*_*A*_ (*green*) fluctuates around a low but constant mean, independently of the Up and Down phases of the neurons that strongly condition the neuronal synaptic variables *s*_*E*_ (*red*) and *s*_*I*_ (*blue*). Of course, the reason why the astrocytic release events are weakly synchronized along the Up and Down phases contrarily to the spiking activity of the neuronal populations is the difference of timescales for information transmission in those cells: on the order of millseconds for neurons versus seconds for astrocytes (the 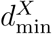 and 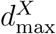 of table 2).

Therefore, in the simulations of figure 5, astrocytes provide the neurons with a constant, basal level of gliotransmission events that fuels their spontaneous collective alternation between Up and Down firing phases. Nevertheless, this background stochastic level of astrocytic input to the neurons is more than an additional random external input to the neurons. To show this, we went back to the spiking model without astrocytes of figure 4 and increased the random external input to the neurons. Figure 4 showed that the network is silent with the default value of the standard deviation of the random external input noise, *σ*_*X*_= 3 mV (table 2). Increasing *σ*_*X*_ to 5 mV does give rise to an Up-Down regime with spontaneous alternation of Up and Down phases (fig. 6). However, the difference between the resulting Up and Down phases is much less marked than in the Up-Down regimes with astrocytes: the subliminal individual membrane voltages are nomore clearly bimodal (fig. 6a), and the difference between firing rates in Up and Down phases is much lower, with a significant firing activity during Down phases (fig. 6b,c).

**Figure 6:**
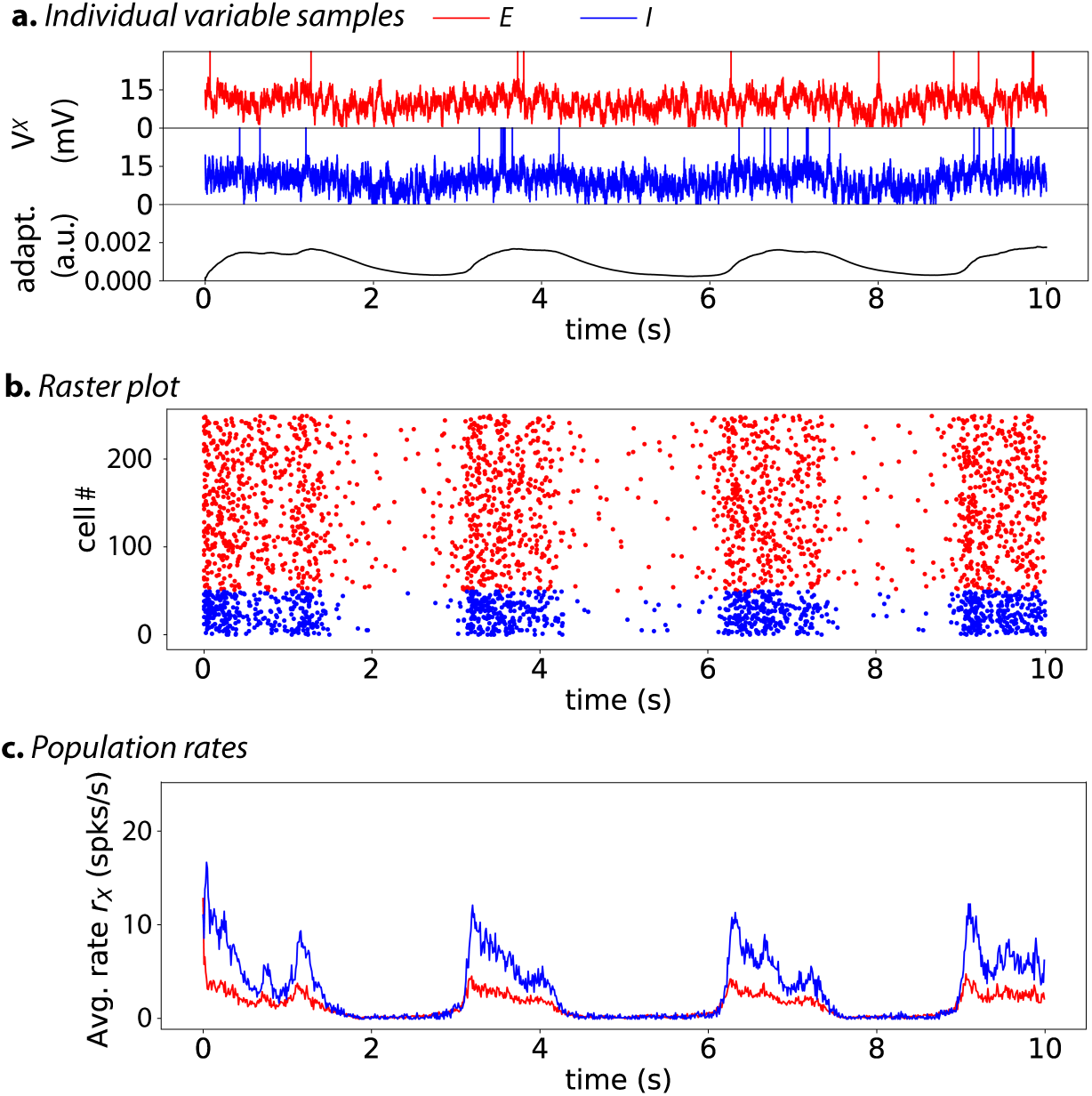
Without astrocytes (*J*_*AE*_ = *J*_*AI*_ = 0, *J*_*IA*_ = *J*_*EA*_ =0 mV) but with large amplitude of the stochastic external input (*a*_*X*_ =5 mV), the stochastic spiking network of sec. 2.2 exhibits spontaneous transitions between Up and Down states, i.e. an Up-Down regime. However the difference between the phases is less marked than the dynamics observed with astrocytes. The membrane voltages of two randomly chosen cells, one excitatory (*red*) neuron, one inhibitory (*blue*) neuron, as well as the average AHP current (*black*) are shown in **(a)**. The spike rastergram **(b)** and the corresponding mean population rates **(c)** are shown using the same color-code. All parameters are identical to those of fig. 4, except for the amplitude of the noise to the neurons *σ*_*E*_ = *σ*_*I*_ =5 mV. *N*_*E*_ = 4,000 excitatory neurons, *N*_*I*_ = 1,000 inhibitory neurons.

Moreover, the range of external input amplitudes that give rise to Up-Down regimes without astrocytes is much more narrow than with astrocytes. Mean-field fixed-point and linear stability analysis of the stochastic spiking network model is shown in Figure 7 (see S1 Text for details). Two bifurcation diagrams are compared: in fig. 7a, astrocytes are absent, like in the simulations of fig. 4, whereas fig. 7b shows the same diagram when astrocytes are present, like in fig. 5. These bifurcation diagrams show the evolution of the fixed points and their stability when one varies the amplitude of the external noisy input, i.e. the standard deviation of the stochastic input to the E and I neurons, *σ*_*X*_. Without astrocytes, the diagram shows a stable fixed-point corresponding to a low firing rate for low *σ*_*X*_ values and a second stable fixed-point yielding a larger firing rate at large *σ*_*X*_ values. In a narrow range of *σ*_*X*_ values ([4.4, 4.5] mV), the two stable fixed points co-exist, together with a third intermediate unstable one (dashed line), thus evidencing a region of bistable dynamics (magnified in the inset). We also indicate with a gray-shaded region the parameter range where simulations of the spiking model evidence spontaneous transitions between Up and Down phases (like in fig. 6). The prediction of the mean-field analysis is not very precise regarding the location of the bistability region, which is probably a finite-size effect related to the finite number of neurons and astrocytes in the simulations. However, the theoretical analysis agrees very well with the narrowness of the bistability region observed in simulations, which confirmes that noise-induced Up-Down regimes with the parameter values of table 2 are observed only for a limited range of input intensities in the absence of astrocytes. In particular, this range is very far from the value *σ*_*X*_ = 3 mV used in the previous simulations, explaining the silent state fig. 4.

**Figure 7:**
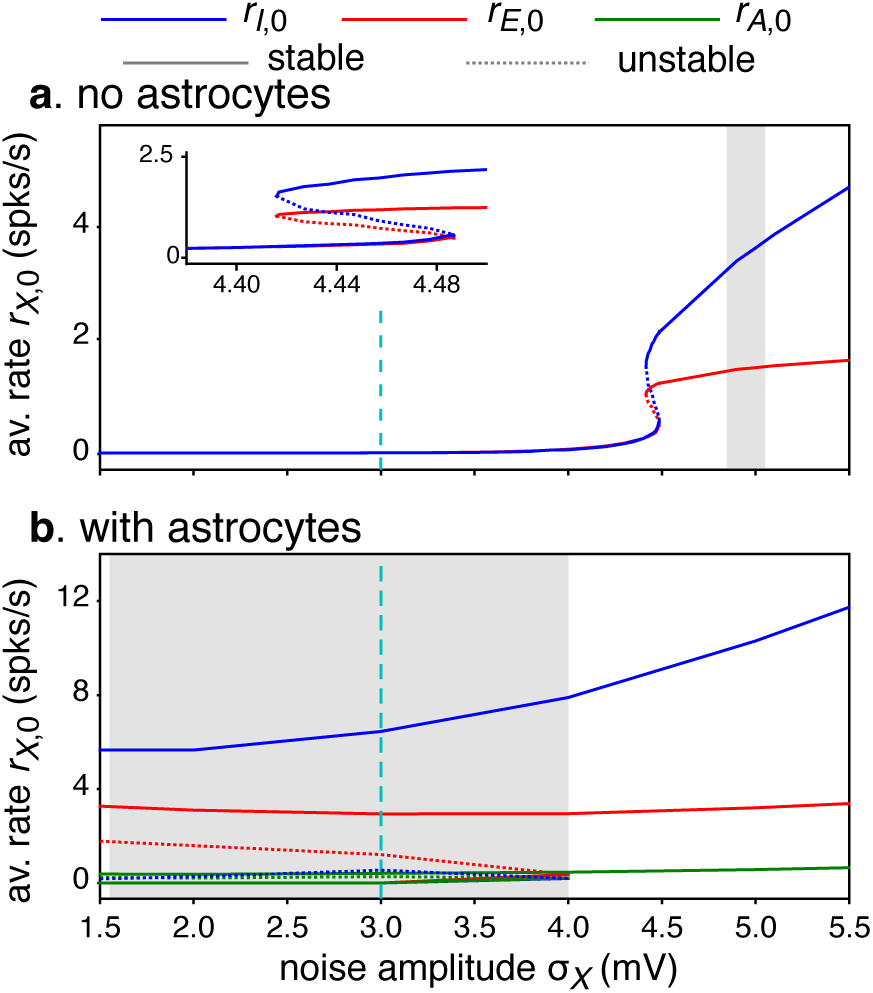
Linear stability analysis of the spiking network model without **(a)** or with astrocytes **(b)** along the intensity of the noisy external input to the neurons *σ*_*X*_. In both cases, a bistable region is observed, ended by a saddle-node bifurcation for large *σ*_*X*_. However, the bistable region is drastically reduced in the absence of astrocytes, as evidenced by the width of the grayshaded region, that locates the range of *σ*_*X*_ values for which Up-Down regimes are observed in numerical simulations of the network. These bifurcation diagrams show the evolution of the stable (solid lines) and unstable (dotted lines) fixed points of the equilibrium rates *r*_*E*,0_ (*red*), *r*_*I*,0_ (*blue*) and *r*_*A*,0_ (*green*). In (a), the insets show a zoom out around the bistability region without astrocytes. See section B of S1 Text for details on linear stability analysis. The dashed cyan vertical line indicates the *β* value used for numerical simulations in fig. 4 and fig. 5. Other parameters are given in table 2. Note in particular that the diagram was obtained using *σ*_*E*_ = *σ*_*I*_ ≡ *σ*_*X*_ and keeping a constant *σ*_*A*_ = 3.

Astrocytes modify part of the bifurcation diagram (fig. 7b): the values of the firing rates of the neurons in the Up and Down stable fixed-point are roughly the same as without astrocytes, but the range of *σ*_*X*_ values for which bistability and Up-Down regimes are observed is incomparably larger, extending to much lower values. In particular, the bistability region now includes the value *σ*_*X*_ = 3 mV, thus the Up-Down regime observed in fig. 4. Note also that the bifurcation analysis predicts that the rate of emission of gliotransmission events by the astrocytes should be very similar either in the Up or the Down state, in strong opposition to the neuronal firing rates. This explains the observation that the gliotransmission emission rate in fig. 5 did not vary much between Up and Down phases: all variables do follow the bistable dynamics of the whole system, but the stable fixed-points for the astrocytes are much closer to each other than those for the neuronal firing rates.

Hence, as for the firing rate model studied in section 3.1 above, adding astrocytes in the spiking model does not drastically alter the nature or number of bifurcations, but relocates the bistability region in the parameter space so that a point in the parameter space that is out of the bistability region without astrocytes can find itself inside the bistability region by the addition of astrocytes, thus exhibiting Up-Down regime.

As a final remark, we note that all the above results were obtained with *J*_*AI*_ > 0, i.e. a scenario where the firing activity of the inhibitory neurons directly increases the probability of gliotransmitter release by the astrocytes. However, we checked that the absence of this specific interaction does not jeopardize the validity of our conclusions here. We show in S1 Figure the results obtained with the rate model (fig. S1a) or the spiking network model (fig. S1b) when the strength of I → A interactions vanishes (*J*_*AI*_ = 0), while keeping all the other parameters as in table 1 or table 2. This figure evidences a handful of changes compared to the scenario *J*_*AI*_ > 0 illustrated above, but the simulation results are still very similar (compare fig. S1a with fig. 3 with or fig. S1b with fig. 5), so that the conclusions drawn above are still valid in the absence of I → A interactions.

## Discussion

Up-Down cortical dynamics have primarily been observed during sleep or anesthesia. However, similar dynamical regimes have also been reported in the cortex during quiet wakefulness (Petersen et al., 2003; Luczak et al., 2007) or during a task (Sachidhanandam et al., 2013; Engel et al., 2016). It thus seems likely that making sense of these dynamics is important for our understanding of brain operations in general, not only during sleep or anesthesia. The cellular mechanisms that support the emergence of spontaneous Up to Down and Down to Up transitions in the cortex are however still unclear. The hypothesis that these transitions could be controlled by a mechanism intrinsic to the neurons of the considered cortical region has been explored by a number of theoretical or computational studies (Bazhenov et al., 2002; Compte et al., 2003; Hill and Tononi, 2005; Benita et al., 2012). However recent experimental studies reported the implication of other types of intrinsic brain cells, in particular astrocytes (Poskanzer and Yuste, 2016, 2011; Sanchez-Vives and McCormick, 2000). These results motivated us to include the contribution of astrocytes in a model of bistable neural network dynamics where Up-Down dynamical regimes could be replicated.

In our numerical simulations, the addition of gliotransmission from astrocytes was indeed sufficient to transform a neural network prepared in the Down, silent state into a dynamical regime of spontaneous alternations between Up and Down states. This result matches in vivo experiments where the activation of a local population of astrocytes was reported to correlate with a transition of the corresponding neural network into the Up-Down regime (Poskanzer and Yuste, 2016). Additional comparisons can be made with the in vivo experiments reported in Jercog et al. (2017) from multichannel silicon microelectrode recordings in the somatosensory cortex of urethane-anesthetized rats. The distributions of Up or Down phase duration in these experiments (their figure 2A) are broad, with Down phases lasting from less than 100 ms to 1.5 s and Up phases reaching larger maximal values, up to 2 s. Our simulation results exhibit similar broad distributions, at least for Up states, a consequence of the large variability of the Up state durations (fig. 5d). The coefficient of variations from the in vivo experiments of Jercog et al. (2017) were 0.61 and 0.70, for Up and Down phases, respectively, to be compared with 0.56 and 0.73 for our simulations. The mean values of the phase durations are also very well replicated by our simulations: 1.03 and 0.46 s for Up and Down phases, respectively, vs 0.65 and 0.38 in the experiment of Jercog et al. (2017), figure 2A. The instantaneous population rate during Up phases in these in vivo experiments is around 4 to 6 Hz in Jercog et al. (2017) (their Figure 1C), a value that is similar to the population rate of excitatory neurons in our simulations (fig. 5d). Taken together, we thus conclude from those quantitative comparisons that our simulation results exhibit Up and Down phases that agree well with available experimental data.

Experimental reports indicate that astrocytes form roughly 20 to 40 % of all glial cells (Verkhratsky and Nedergaard, 2018). On the other hand, estimates of the ratio between glial cells and neurons in human cortex varies from 1.5 to more than 2 in humans (Verkhratsky and Nedergaard, 2018). Altogether, those numbers yield an astrocyte:neuron number ratio in the human cortex that ranges from 1:3 to 1:1. The numbers used in our simulations of the spiking network model are in good agreements with these experimental reports, with an astrocyte:neuron number ratio of 1:2.5. In the present article, we adopted the modelling choice made by Jercog et al. (2017) for their spiking model where the synapse dynamics are modelled using a single population variable, integrating the spikes emitted by the whole population into a single variable that can then be fedback to the other cells. This choice limits the range of modelling exploration regarding connectivity. It forbids models where the inputs received by an astrocyte is restricted to a subset of the neurons or, conversely, those where gliotransmission from an astrocyte targets only a subset of the synapses of a neuron. On the other hand, though, this modelling choice greatly facilitates theoretical (mean-field) analysis of the stochastic network model. We believe that the possibility to rely simulation results on an underlying sound theoretical analysis was important for the present article, and this is the reason why we have chosen to keep these population synapses. We leave for future works the study of models that would incorporate the main ingredients of our models above, but with real individual synapses.

Another simplification we made in the model was to use mathematical expressions for astrocyte gliotransmitter release rate or gliotransmission events that were borrowed from classical models for neuronal firing rate or membrane potential dynamics. We acknowledge that using equation (5) or equation (9) is a strongly simplified modeling of gliotransmission. However, it has the advantage of preserving the main biological ingredients of gliotransmission while keeping the model simple enough that we can still proceed to some analytical study of its stability. Moreover, we think that our choice of parameters within this modelling frameworks made it possible to capture at least part of the fundamental differences of signal integration in neurons versus astrocytes. In particular, the signalling delay in our spiking network model was kept three orders of magnitude larger in astrocytes compared to neurons, i.e. seconds versus milliseconds. This difference of timescales turned out to be crucial for the network dynamics illustrated in fig. 5 where a stationary background of astrocytic gliotransmission events triggers spontaneous transitions between synchronized Up and Down phases of neuronal firing. Another major conclusion from our study is that gliotransmission in the model does not have a drastic effect on the underlying dynamics of the network. Adding gliotransmission does not modify the number nor the type of the observed bifurcations, it only alters the values of the parameters at which these bifurcations occur. As a result, glio-transmission can transform a silent neural network model into a network exhibiting Up-Down dynamics, with no change of the neuron-related parameters, and no alterations of the neural mechanisms that control the transitions between Up and Down phases. The modeling literature proposes mathematical descriptions of the process of gliotransmitter release from astrocytes that are much more complex or accurate than the simple phenomenological expressions used here, see e.g., De Pitta and Berry (2019) for a recent account. However the price to pay for the added complexity would be a restriction of the available mathematical understanding of the system dynamics. Future numerical simulation works will be needed to assess whether the inclusion of such more complex descriptions comes with important changes of the main conclusions of the present study.

The main experimentally-testable prediction made by our work is arguably the possibility of a dynamical regime where the astrocytic gliotransmitter release events are only weakly synchronized to the succession of Up and Down phases of the neuron firing state, i.e. the population frequency of gliotransmitter release events does not change much in Up phases compared to Down phases. Experimental testing of this prediction would consist in measuring simultaneously the activity of a local population of neurons using e.g., multi-channel silicon microelectrodes while monitoring the gliotransmitter events from astrocytes from the same local area. Gliotransmitter release events are difficult to monitor experimentally, even with glutamate-sensitive fluorescent reporters (see e.g., fig. 7D in Poskanzer and Yuste (2016)). Monitoring intracellular calcium activity could constitute a good proxy to locate glutamate release events by astrocytes. However, recent experimental studies have challenged the relation between calcium signals recorded from astrocyte cell bodies from those initiated in the fine processes, that are expected to contact the synapses (Rusakov, 2015; Shigetomi et al., 2016; Bindocci et al., 2017). Therefore, experimental testing of the above dynamical regime would probably need the measure of local calcium signals, within the fine astrocyte processes that form the so-called ”gliapil”. At any rate, this predicted dynamical regime is supported by activity-dependent release of gliotransmitters by astrocytes, which existence and impact on the neurons in physiological conditions is still debated among experimental neuroscientists (see e.g. Savtchouk and Volterra, 2018; Fiacco and McCarthy, 2018). Therefore, according to the work presented here, experimental observation of astrocytes releasing gliotransmitters at a roughly constant rate while neurons undergo successive Up and Down firing phases, should be interpreted as an argument in favor of the existence of gliotransmission, and not against it.

