## Supporting Information for "Modelling the modulation of cortical Up-Down state switching by astrocytes"

[1] Inria, 69100 Villeurbanne, France.

[2] University of Lyon, LIRIS UMR5205, 69100 Villeurbanne, France.

#### Supporting Information

#### S1 Text. Fixed points and linear stability analyses

##### A. Rate Model

**Noiseless model**. We start with the rate model defined by equations (1) to (6) and first neglect the external noisy input. In this case, the nullclines of the system are given by

$$r_E = g_E[I_E - a - \theta_E]_+ \tag{S1.1}$$

$$r_I = g_I[I_I - \theta_I]_+ \tag{S1.2}$$

$$r_A = g_A[I_A - \theta_A]_+ \tag{S1.3}$$

$$a = \beta r_E \tag{S1.4}$$

Note that because of the rectification functions in equations (S1.1) to (S1.3), the values of the rates at a fixed point cannot be negative. The rate model equations (1) to (6) being a piecewise-smooth system, a rigorous analysis of the stability of its fixed points would require dedicated analysis methods (see e.g., di Bernardo et al., 2008). Here, we leave this analysis for further work and assume that all fixed points remain far from the switching manifolds where the arguments of the rectification functions change signs and proceed to linear stability analysis in each of the respective regions.

A Down or silent state can be characterized as a fixed-point where both neuronal populations are silent, i.e. $r_E = r_I = 0$. Equation (S1.4) means that adaptation $a$ also vanishes in such a Down state. The rectification functions of equations (S1.1) and (S1.2) impose $\theta_E \geq 0$ and $\theta_I \geq 0$ for the Down fixed-point to exist. Indeed $\theta_E < 0$ would mean from eq. (S1.1) that $r_A < 0$ at the fixed-point (since $r_E = 0$), which is not compatible with eq. (S1.3). Hence the Down state exists only if $\theta_E \geq 0$ or $\theta_I \geq 0$. If, in addition $\theta_A \geq 0$, the Down state is $(r_E, r_I, r_A, a) = (0, 0, 0, 0)$. Assuming that all the rectification functions in eq. (1), eq. (2) and eq. (5) vanish in the neighborhood of the fixed point, we find that this Down state is stable by linear stability analysis (eigenvalues of the Jacobian:$\{-1/\tau_E, -1/\tau_I, -1/\tau_A, -1/\tau_a\}$).

In the case $\theta_A < 0$ (still with $\theta_I \geq 0$ and $\theta_E \geq 0$), we assume that the argument of the rectification function in eq. (5) is strictly positive, while the rectification functions in equations (1) and (2) vanish. The nullcline for $r_A$, eq. (S1.3) then becomes $r_A = g_A(J_{AA} r_A - \theta_A)$. Therefore, there still is a positive Down fixed-point $(r_E, r_I, r_A, a) = (0, 0, -g_A\theta_A/(1 - g_A J_{AA}), 0)$ but only for:

$$\theta_E > -g_A J_{EA} \theta_A/(1 - g_A J_{AA}), \tag{S1.5}$$

as well as $\theta_I > -g_A J_{IA} \theta_A/(1 - g_A J_{AA})$ and $g_A J_{AA} < 1$. Close to this fixed-point, the Jacobian matrix reads

$$\begin{pmatrix} -1/\tau_E & 0 & 0 & 0 \\ 0 & -1/\tau_I & 0 & 0 \\ 0 & 0 & \left(g_A J_{AA} - 1\right)/\tau_A & 0 \\ \beta & 0 & 0 & -1/\tau_a \end{pmatrix} \tag{S1.6}$$

so stability is granted whenever $g_A J_{AA} < 1$, i.e as soon as the equilibrium value for $r_A$ exists.

To find an Up state fixed-point with non-zero rates we follow Jercog et al. (2017) and substitute the value of the adaptation at equilibrium $a = \beta r_E$, assuming that the arguments of all the rectification functions of eq. (1), eq. (2) and eq. (5) are strictly positive. This yields:

$$r_E = \frac{h_E\theta_E + f_{EI}\theta_I + f_{EA}\theta_A}{|M|}, \tag{S1.7}$$

$$r_I = \frac{f_{IE}\theta_E + (h_I - \beta J'_{AA})\theta_I + (f_{IA} + J_{IA}\beta)\theta_A}{|M|} \tag{S1.8}$$

and

$$r_A = \frac{f_{AE}\theta_E + (f_{AI} + J_{AI}\beta)\theta_I + (h_A - \beta J'_{II})\theta_A}{|M|} \tag{S1.9}$$

with

$$J'_{XY} = J_{XY} - \frac{1}{g_X}, \tag{S1.10}$$

$$h_X = J'_{YY}J'_{ZZ} - J_{YZ}J_{ZY}, \tag{S1.11}$$

$$f_{XY} = J_{XZ}J_{ZY} - J_{XY}J'_{ZZ}, \tag{S1.12}$$

and

$$|M| = J_{AE}f_{EA} + J_{AI}(f_{IA} + J_{IA}\beta) + J'_{AA}(h_A - \beta J'_{II}) \tag{S1.13}$$

One condition for the UP state fixed-point to exist is that the right hand side of eq. (S1.7), eq. (S1.8) and eq. (S1.9) are positive. Given our reference parameters (table 1), the condition on $r_I$, i.e., $r_I > 0$ is the most restrictive condition. In other words, when $\theta_E$ and $\beta$ are varied, eq. (S1.8) is the first one to become positive. Moreover, with our reference parameters, it turns out that the determinant $|M| < 0$. The condition for the existence of the Up state fixed-point thus becomes

$$\beta < \frac{f_{IE}\theta_E + h_I\theta_I + f_{IA}\theta_A}{J'_{AA}\theta_I - J_{IA}\theta_A}. \tag{S1.14}$$

since $J'_{AA}\theta_I - J_{IA}\theta_A < 0$ with our reference parameters (table 1).

Considering that in the neighborhood of the Up fixed-point, all the rectification functions of the model are positive, the Jacobian matrix reads

$$\begin{pmatrix} \frac{-1+g_E(J_{EE}-\beta)}{\tau_E} & \frac{g_E J_{EI}}{\tau_E} & \frac{g_E J_{EA}}{\tau_E} \\ \frac{g_I J_{IE}}{\tau_I} & \frac{-1+g_I J_{II}}{\tau_I} & \frac{g_I J_{IA}}{\tau_I} \\ \frac{g_A J_{AE}}{\tau_A} & \frac{g_A J_{AI}}{\tau_A} & \frac{-1+g_A J_{AA}}{\tau_A} \end{pmatrix}.$$

It is possible to obtain analytical expressions for the eigenvalues of this matrix, however these expressions are too complex to be really useful, even to determine their signs. We therefore estimated their values numerically to explore stability of the Up fixed-point, with numerical estimation of the sign of the real part of the eigenvalues of the matrix. Note finally that in regions where both the Up state and the Down state fixed-points are stable, the existence of a third, unstable and intermediate fixed-point can be evidenced by numerical analysis.

**The effect of noise on the model**. The addition of noise however complicates the above picture. In particular, spontaneous Up-Down transitions can occur even in the region where the Down state is stable and the Up state unstable, through a dynamical regime whereby noise triggers the Down to Up switches and adaptation triggers the reverse Up to Down transitions. In Jercog et al. (2017), it is proposed that this subregion can be estimated to start when the Up state is unstable under the sole influence of adaptation, a condition that can be deduced from eq. (S1.14) with $\beta = 0$:

$$\theta_E < -\frac{h_I \theta_I + f_{IA} \theta_A}{f_{IE}} \tag{S1.15}$$

Likewise, the symmetrical regime exists in the region where the Up state is stable and the Down state unstable, for which Up-to-Down transitions are triggered by noise and Down-to-Up switches by adaptation. Jercog et al. (2017) proposes to delineate this region by the situation where adaptation in the Up-state is large enough to counterbalance the effect of $\theta_E$, i.e. $\beta r_E + \theta_E > 0$, where $r_E$ is given by eq. (S1.7), which yields

$$\beta > \frac{-\theta_E (J_{AE} f_{EA} + J_{AI} f_{IA} + J'_{AA} h_A)}{f_{EI} \theta_I + f_{EA} \theta_A + (h_E - J'_{AA} J'_{II} - J_{AI} J_{IA}) \theta_E} \tag{S1.16}$$

##### B. Spiking Model

Analysis of the spiking model defined by equations (7) to (13) was carried out as follows. The nullclines of the population averaged rates are obtained from the equilibrium firing rate ($r_{X,0}$), which is given by the self consistent mean-field equation (Brunel and Hakim, 1999):

$$r_{X,0} = \frac{1}{\tau_X}\left[\int_0^\infty \frac{dy}{y} e^{-y^2}(e^{2yy_t^X} - e^{2yy_r^X})\right]^{-1} \quad \text{(S1.17)}$$

with

$$y_r^X = \frac{V_r - I_{X,0}}{\sigma_X} \qquad y_t^X = \frac{\theta_X - I_{X,0}}{\sigma_X} \quad \text{(S1.18)}$$

and with the currents:

$$I_{E,0} = V_{L,E} + I_{rec,E} - K_a \beta r_{E,0} \quad \text{(S1.19)}$$

$$I_{I,0} = V_{L,I} + I_{rec,I} \quad \text{(S1.20)}$$

$$I_{A,0} = V_{L,A} + I_{rec,A} \quad \text{(S1.21)}$$

$$I_{rec,X} = C_{XE}J_{XE}r_{E,0}\underline{\tau_E} + C_{XI}J_{XI}r_{I,0}\underline{\tau_I} + C_{XA}J_{XA}r_{A,0}\underline{\tau_A} \quad \text{(S1.22)}$$

where $C_{XY}$ is the connectivity (0.1 for astrocyte to neuron connection, 0.5 for neuron to astrocytes, 1.0 otherwise). Note that we replaced $I_{a,0} = \beta r_{E,0}$. These self-consistent equations are solved by finding the intersection between the right and the left side of equation (S1.17). The fixed points are the intersections of the three surfaces $r_{E,0}$, $r_{I,0}$ and $r_{A,0}$. The stability of these fixed points is determined with the sign of the eigenvalue $\lambda$ (Brunel and Hakim, 1999; Roxin and Compte, 2016; Ledoux and Brunel,

2011). Solving the system

$$\begin{cases} \delta r_E = F_{EE}(\lambda)\delta r_E + F_{EI}(\lambda)\delta r_I + F_{EA}(\lambda)\delta r_A \\ \delta r_I = F_{IE}(\lambda)\delta r_E + F_{II}(\lambda)\delta r_I + F_{IA}(\lambda)\delta r_A \\ \delta r_A = F_{AE}(\lambda)\delta r_E + F_{AI}(\lambda)\delta r_I + F_{AA}(\lambda)\delta r_A \end{cases} \tag{S1.23}$$

for the perturbations around the fixed-point, $\delta r_E$, $\delta r_I$ and $\delta r_A$, one gets the condition (omitting the dependences on $\lambda$ for readability):

$$\begin{aligned} &(F_{EE}-1)(F_{II}-1)(F_{AA}-1) + F_{EI}F_{IA}F_{AE} + F_{EA}F_{IE}F_{AI} \\ -&(F_{EE}-1)F_{IA}F_{AI} - F_{EI}F_{IE}(F_{AA}-1) - F_{EA}(F_{II}-1)F_{AE} = 0 \end{aligned} \tag{S1.24}$$

with

$$F_{XY}(\lambda) = J_{XY}R_X(\lambda)S_Y(\lambda) \tag{S1.25}$$

and the synaptic response function $S_Y(\lambda)$

$$S_Y(\lambda) = \frac{e^{-\lambda d^Y}}{(1+\lambda\tau_r^Y)(1+\lambda\tau_d^Y)} \tag{S1.26}$$

In this equation, $R_X(\lambda)$ is the neuronal response function defined as

$$R_X(\lambda) = \frac{r_{X,0}}{\sigma_X(1+\lambda\tau_X)} \frac{\frac{\partial U}{\partial y}(y_t^X, \lambda\tau_X) - \frac{\partial U}{\partial y}(y_r^X, \lambda\tau_X)}{U(y_t^X, \lambda\tau_X) - U(y_r^X, \lambda\tau_X)} \tag{S1.27}$$

with

$$U(y,\lambda)=\frac{e^{y^2}}{\Gamma\left[\frac{1+\lambda}{2}\right]}M\left(\frac{1-\lambda}{2},\frac{1}{2},-y^2\right)+\frac{2ye^{y^2}}{\Gamma\left[\frac{\lambda}{2}\right]}M\left(1-\frac{\lambda}{2},\frac{3}{2},-y^2\right) \tag{S1.28}$$

with $M$ is a confluent hypergeometric function. Stability is assessed by solving eq. (S1.24) for $\lambda$ numerically. The fixed point is stable when its real part is negative.

### S1 Figure. Results in the absence of I → A interactions

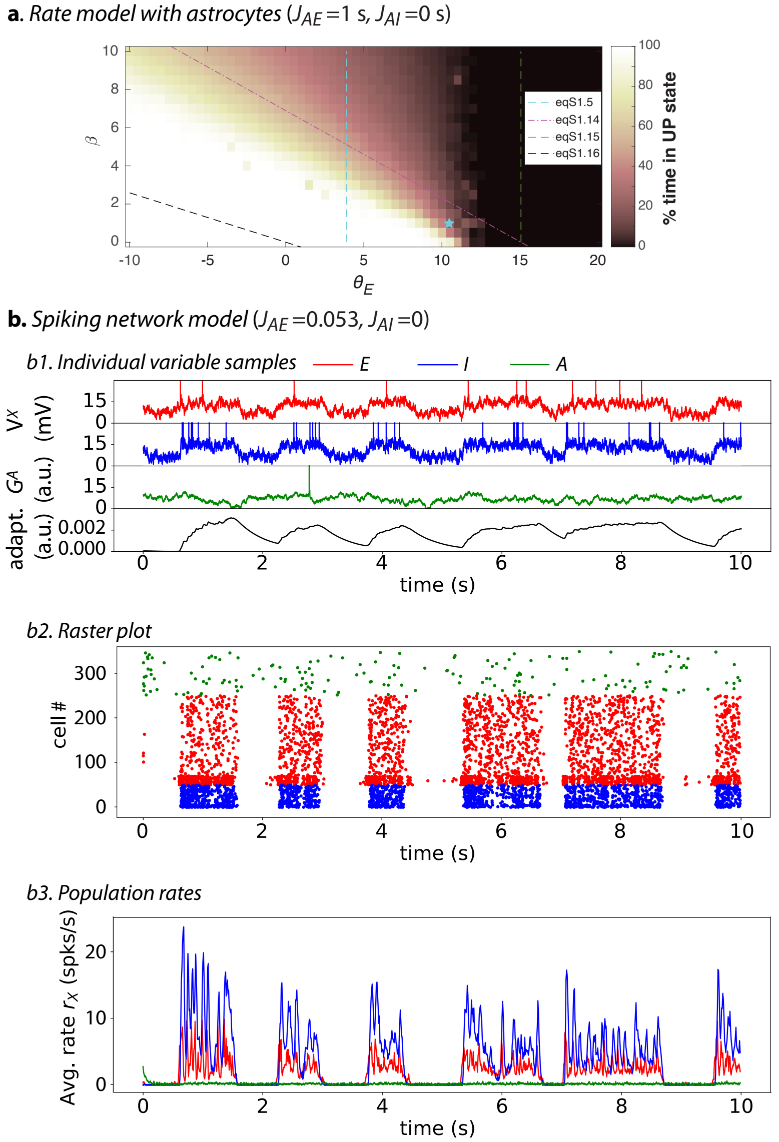


**Figure S1:** Results of the rate model of equations (1) to (6) **(a)** or the spiking network model of equations (7) to (13) **(b)** obtained in the presence of astrocytes, but with $J_{AI} = 0$, i.e. with no direct effect of inhibitory neurons on astrocytic gliotransmitter release. All parameters were as indicated in table 1 or table 2, except for the value of $J_{AI}$ that was set to 0. Refer to fig. 3c and fig. 5 for the color-codes and parameters of panels (a) and (b), respectively.
